# Hippocampus-reuniens beta coupling supports goal-directed spatial navigation

**DOI:** 10.64898/2026.05.20.726518

**Authors:** Tristan Baumann, Hao Mei, Oxana Eschenko

## Abstract

Prefrontal–hippocampal communication, supported by both direct and indirect anatomical pathways, is essential for a range of cognitive functions, including spatial navigation. The midline thalamic nucleus reuniens (RE) has been proposed as a key hub along this indirect pathway, coordinating bidirectional interactions between the hippocampus (HC) and prefrontal cortex (PFC). Here, we investigated functional dynamics within the HC–RE–PFC network as adult male rats learned to navigate a complex maze. Initial behavioral analyses revealed three distinct learning phases: exploration, goal-oriented learning, and efficient navigation. Aligning neural data with subject-specific transitions between these phases uncovered distinct neural signatures associated with each learning phase. Notably, the transition from exploratory to goal-directed behavior was accompanied by the emergence of persistent HC-RE beta-band (15-25Hz) interactions, including elevated beta coherence, theta-beta phase-amplitude coupling, and HC-to-RE Granger causality. The interactions further scaled with navigational efficiency, showing increased HC-RE beta coherence in trials without errors. Together, these findings provide new evidence for dynamic HC–RE interactions during goal-directed navigation and support an emerging view that RE functions as a working memory buffer for route-related information, revealing a potential network mechanism underlying flexible spatial behavior.

**Highlights:**

- Beta-band hippocampus–reuniens coupling emerges during goal-directed navigation.
- PAC between HC theta and RE beta increased during goal learning.
- HC–RE coupling strength increases with navigational efficiency.
- Directional hippocampus-to-reuniens communication emerges during learning
- Reuniens supports hippocampal–prefrontal integration for flexible behavior

**Significance Statement:** Flexible spatial navigation requires coordinated communication between the hippocampus (HC) and the prefrontal cortex (PFC), yet the thalamic mechanisms coordinating HC-PFC interaction remain poorly understood. This study reveals that the nucleus reuniens (RE) of the midline thalamus plays a critical role in mediating HC-PFC communication during spatial learning. By combining detailed behavioral analysis with simultaneous multi-site electrophysiological recording, we identify beta-band HC–RE coupling as a neural signature of the transition from exploration to goal-directed navigation. The strength and directionality of this interaction tracked improvements in navigational efficiency, revealing a thalamic mechanism for integrating hippocampal output into prefrontal circuits. These findings highlight the RE as a working memory buffer for route-related information and advance our understanding of how thalamic hubs contribute to the circuit-level dynamics underlying flexible behavior.

## Introduction

Successful goal-directed spatial navigation relies on coordinated activity across distributed brain networks, with the hippocampus (HC) and prefrontal cortex (PFC) playing a central role (***Granon and Poucet, 1995; Colgin, 2011; Ito et al., 2015; Ito, 2018***). The HC supports rapid encoding of spatial information and stabilization of memory traces (***Preston and Eichenbaum, 2013; Jayachandran et al., 2023***), whereas the PFC contributes contextual representation, selective memory retrieval, and action planning (***Xu and Südhof, 2013; Eichenbaum, 2017***). In rodents, interactions between the dorsal HC and medial PFC are essential for spatial working memory, long-term memory, and decision-making (***Buzsáki, 1996; Colgin, 2011; Eichenbaum, 2017***), and disruptions in HC-PFC communication lead to marked deficits in spatial cognition and memory (***Xu and Südhof, 2013; Lopez et al., 2012; Wirt and Hyman, 2017***).

Spatially and temporally coordinated brain oscillations are thought to be a central mechanism enabling cross-regional communication. These rhythmic patterns, arising from synchronous firing of large neuronal populations, are readily detected in local field potentials (LFPs) and electroen-cephalography (EEG) as large-amplitude fluctuations in extracellular currents (***Colgin, 2016***). Neural oscillations provide a temporal framework for controlling the timing of neuronal firing and crossregional coupling, thereby enabling the encoding, transfer, and retrieval of information (***Fries, 2005; Engel and Fries, 2010; Colgin, 2016***).

In rodents, hippocampal theta oscillations (6-11Hz), which prominently appear during locomotion, are thought to play an essential role in HC-PFC communication. Several studies have reported phase-locking of prefrontal neurons to hippocampal theta in animals performing spatial working memory tasks, with the strength of this coupling correlating with working memory accuracy (***Jones and Wilson, 2005; Hyman et al., 2010; Hallock et al., 2016***). Similarly, spatial task performance is associated with increased HC–PFC coherence in both theta (***Jones and Wilson, 2005; Benchenane et al., 2010; Hallock et al., 2016; de Mooij-van Malsen et al., 2023***) and gamma (30–120 Hz) bands (***Spellman et al., 2015; Bygrave et al., 2019; de Mooij-van Malsen et al., 2023***).

These functional interactions occur despite relatively asymmetrical direct anatomical connectivity. In rodents, robust monosynaptic projections extend from ventral CA1 and subiculum to the medial PFC (***Swanson, 1981; Vertes, 2006; Wirt and Hyman, 2017***), whereas direct PFC-to-HC projections are sparse (***Dolleman-van der Weel et al., 2019***). In mice, a monosynaptic pathway from the anterior cingulate cortex to CA1 and CA3 has been described (***Rajasethupathy et al., 2015***), but overall, HC receives limited direct inputs from PFC. The lack of strong reciprocal connectivity points to the importance of indirect pathways, for example, via the thalamus.

The midline thalamic nucleus reuniens (RE) has been proposed as a key nucleus along this indirect pathway, coordinating HC-PFC communication (***Cassel et al., 2013, 2021; McKenna and Vertes, 2004; Scheel et al., 2020; Vertes et al., 2006; Dolleman-van der Weel et al., 2019***). RE receives input from deep layers of the prelimbic and infralimbic cortices and projects back to both superficial and deep layers (***Scheel et al., 2020; McKenna and Vertes, 2004; Vertes et al., 2006***). It is also densely innervated by the ventral HC and subiculum and projects to the stratum lacunosum-moleculare of the ventral HC, targeting distal dendrites of excitatory and inhibitory neurons (***Scheel et al., 2020; Varela et al., 2014; Vertes et al., 2015***). Optogenetic activation of RE evokes excitatory responses in CA1 (***Hauer et al., 2019***), and a subset of RE neurons sends collaterals to both HC and PFC (***Hoover and Vertes, 2012; Varela et al., 2014***), providing a potential mechanism for simultaneous modulation of both regions.

Functionally, RE has been implicated in multiple memory processes. Single units in RE show trajectory-dependent coding during spatial working memory tasks (***Ito et al., 2015***). Lesions or inactivation of RE lead to deficits in spatial working memory (***Davoodi et al., 2009; Hallock et al., 2016; Hembrook and Mair, 2011; Hembrook et al., 2012; Layfield et al., 2015; Maisson et al., 2018; Viena et al., 2018***), spatial memory consolidation (***Loureiro et al., 2012; Xu and Südhof, 2013***), and retrieval (***Mei et al., 2018; Barker and Warburton, 2018***). Inactivation of RE also reduces HC-PFC oscillatory synchrony across a wide range of frequencies, including theta, beta, and gamma during wakefulness (***Kafetzopoulos et al., 2018; Totty et al., 2023***), and slow oscillations (∼1 Hz) and gamma during sleep (***Ferraris et al., 2018; Hauer et al., 2019***).

These findings support the view that RE plays an important role in coordinating the extended neural network underlying spatial cognition and working memory. However, the specific function, temporal dynamics, and behavioral correlates of RE-mediated HC–PFC interactions remain incompletely understood.

Here, we investigated functional interactions within the HC-RE-PFC network while rats learned to efficiently navigate a complex maze to obtain rewards. Due to the established role of RE in spatial cognition, including results from an earlier pharmacological study using the same behavioral paradigm (***Mei et al., 2018***), we hypothesized that spatial learning would be accompanied by dynamic changes in coordinated HC-RE-PFC activity. Based on behavior, we identified subject-specific transitions between distinct learning phases, each associated with distinct neural signatures. Notably, we found persistent HC-to-RE beta-band (15–25 Hz) interactions, which emerged when animals transitioned from exploratory to goal-oriented behavior, with interaction strength increasing alongside navigational efficiency. The results suggest that HC–RE beta coupling in particular may support goal-directed spatial navigation.

## Results

Six adult male rats were chronically implanted with electrodes targeting the medial PFC, RE, and dorsal HC (***Figure 1*** a). The rats were trained on a spatial navigation task in an elevated maze with a crossword-like layout containing multiple intersections and dead ends (***Figure 1*** b). Two salient visual cues were affixed to a curtain surrounding the maze to provide an extra-maze spatial reference. The rats were accustomed to the experimental environment, but the maze layout and reward location were novel at the start of training.

**Figure 1.**
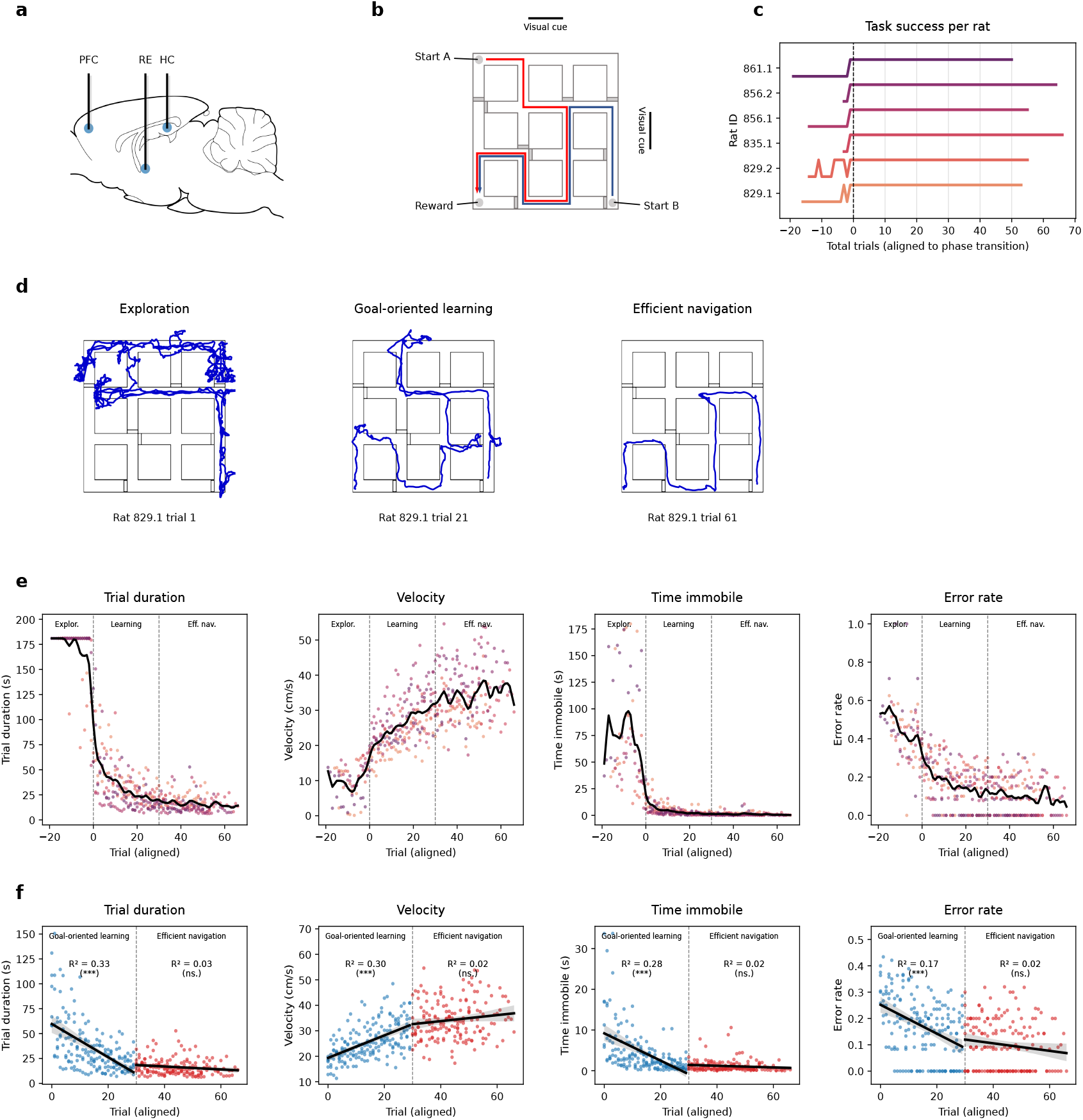
Experimental setup and behavioral dynamics during spatial learning. **(a)** Electrode placement in the rat brain, targeting medial PFC, RE, and dorsal HC. Schematic based on ***Paxinos and Watson*** (***2006***). **(b)** Crossword maze layout indicating the shortest routes from start to goal. Gray boxes indicate blocked-off arms. Two large visual cues were affixed to the curtain surrounding the maze to serve as an extra-maze reference. The maze did not contain any loops. **(c)** Probability of reaching reward, aligned to the last unsuccessful trial. A trial was counted as unsuccessful if the animal failed to reach the reward within 3 minutes. Learning was split into three phases based on this alignment. Trials <0 were assigned to exploration, trials 0-29 to goal-oriented learning, and trials ≥30 to efficient navigation. **(d)** Representative trajectories for each learning phase. Navigation in the exploration phase was characterized by slow, circuitous movements, frequent pauses, and a failure to reach the reward. Trajectories during goal-oriented learning always terminated at the reward, but often contained navigational errors. In the efficient navigation phase, rats made very few navigational errors and quickly reached the reward by following the shortest path. **(e**,**f)** Behavioral dynamics of spatial learning. **e**, Behavioral measures were aligned to the subject-specific transition point from exploration to goal-oriented behavior (e) and the beginning of the goal-directed phase (f). Color-coded dots show trial averages for individual rats; the black lines show the grand average. Note, the transition from exploration to goal-oriented behavior was characterized by a step-like change in most behavioral measures, while goal-oriented learning was associated with a slower dynamics. Linear regression lines (f) highlight gradual behavioral optimization during goal-oriented learning (blue) and asymptotic performance during efficient navigation (red).

Over seven daily sessions of 10 trials each, the rats learned to navigate the maze efficiently to retrieve a food reward. At the start of each trial, the animals were released from one of two pseudorandomized starting points and given up to three minutes to reach the reward port. By providing allocentric cues and varying the starting location, the task was intended to promote HC-dependent navigation and HC-PFC interaction. To minimize procedural learning, training was stopped after seven sessions (70 trials), which was sufficient to reach an asymptotic performance level. Local field potentials (LFPs, 1-500Hz) were recorded simultaneously from the medial PFC, RE, and dorsal HC throughout the task.

### Behavioral switch from spatial exploration to goal-directed navigation

During early trials, behavior was variable and often included prolonged pauses, grooming, rearing, or stationary periods. Rats required between 3 and 19 trials to reach the reward port for the first time. Following the first successful trial, 4 out of 6 animals consistently located the reward on all subsequent trials. The remaining two required an additional 2 and 10 trials, respectively (***Figure 1*** c). Example trajectories across learning are shown in ***Figure 1*** d.

We quantified performance using four behavioral measures: trial duration, mean velocity, time spent immobile (velocity < 5cm/s), and path efficiency. Path efficiency was estimated as the number of deviations from the shortest possible route, including incorrect turns and backtracking, normalized by the number of decision points encountered to yield a trial-wise error rate.

Notably, all measures showed a sharp, step-like change once the rats began reaching the reward reliably: trial duration and time immobile decreased abruptly, while velocity and path efficiency increased (***Figure 1*** d). After this initial transition, performance continued to improve gradually, approaching an asymptote after approximately 30 additional trials. Based on these patterns, we defined three learning phases:

1. **Exploration**, the trials before consistent reward retrieval (range: 3-19 trials)
2. **Goal-oriented learning**, the next 30 trials following the switch
3. **Efficient navigation**, the remaining trials (range: 21-37 trials)

We next compared each behavioral metric across learning phases, factoring in the starting points. A two-way repeated-measures ANOVA (phase × start, FDR-corrected) revealed a strong main effect of learning phase on all metrics (trial duration: *F* (2, 10) = 553.32, *p* < 0.001, 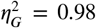; velocity: *F* (2, 10) = 98.09, *p* < 0.001, 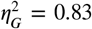; immobility: *F* (2, 10) = 20.94, *p* < 0.001, 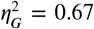; path efficiency: *F* (2, 10) = 105.39, *p* < 0.001, 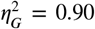). There were no main effects of starting point (all *F* (1, 5) < 3.58, *p* > 0.174, 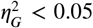) and no significant interactions (all *F* (2, 10) < 4.23, *p* > 0.111, 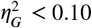). Data from both starting points were therefore pooled for subsequent analyses. Post-hoc pairwise comparisons are summarized in ***Table 1***.

**Table 1.** Behavior by learning phase. Significant differences between successive task phases (goal-oriented learning vs. exploration and efficient navigation vs. goal-oriented learning) are highlighted in bold and marked by asterisks (*p* < 0.01: **; *p* < 0.001 ***, post-hoc paired t-test).

|  | Exploration<br>(3-19 trials) | Goal-oriented learning<br>(30 trials) | Efficient navigation<br>(21-37 trials) |
| --- | --- | --- | --- |
| Trial duration (s) | 169.68 ± 3.52 | **35.44 ± 1.90 (\*\*\*)** | **16.32 ± 0.63 (\*\*\*)** |
| Velocity (cm/s) | 10.05 ± 0.58 | **25.76 ± 0.52 (\*\*\*)** | **34.28 ± 0.58 (\*\*\*)** |
| Time immobile (s) | 71.22 ± 5.80 | **4.32 ± 0.41 (\*\*)** | **1.12 ± 0.11 (\*\*)** |
| Error rate | 0.45 ± 0.02 | **0.17 ± 0.01 (\*\*\*)** | **0.10 ± 0.01 (\*\*)** |

Fitting linear regression models to trial-by-trial behavioral data confirmed distinct early and late learning phases (***Figure 1*** e). All behavioral metrics showed steeper slopes in the goal-oriented learning phase (all *R*^2^ > 0.17, *p* < 0.001), whereas slopes during the efficient navigation phase were minimal and non-significant (all *R*^2^ < 0.03, *p* > 0.050).

In summary, behavioral analysis identified three distinct learning phases: Exploration, goal-oriented learning, and efficient navigation. The exploration phase was characterized by slow velocity, frequent immobility, and a high error rate. After the first successful trial, animals rapidly shifted to faster and increasingly efficient trajectories characteristic of the goal-oriented learning phase. The task performance eventually approached an asymptote, showing only minor improvements over the last ∼30 trials in the efficient navigation phase. Notably, the transition from exploration to goal-oriented learning was accompanied by both qualitative and quantitative changes in behavior, consistent with a shift in navigational strategy.

### HC–RE–PFC network dynamics reflect a strategy shift

To identify how the learning phases mapped onto neural dynamics, we analyzed LFPs recorded simultaneously from HC, RE, and PFC across all trials. A total of 406 artifact-free trials were included in the analysis (n = 6 rats, average 67.7 trials per rat). Trials were categorized by learning phase, and LFP analyses were performed on pooled data, since we did not observe significant behavioral differences between starting points.

#### LFP Power Spectrum indicates a goal-oriented increase in hippocampal beta

We first examined whether LFP spectral power differed between learning phases. For spectral analysis, each trial was segmented into 1s epochs (0.5s overlap), and power spectral densities were estimated using Welch’s method. The analysis was restricted to locomotor epochs (mean velocity >5cm/s). Across all regions, broadband spectral power showed multiple peaks, primarily in the theta and beta ranges. Based on the peaks and troughs of the power spectra, we defined four frequency bands: theta (6-11 Hz), beta (15-25 Hz), low gamma (30-50 Hz), and high gamma (50-100 Hz) (***Figure 2*** a,b).

**Figure 2.**
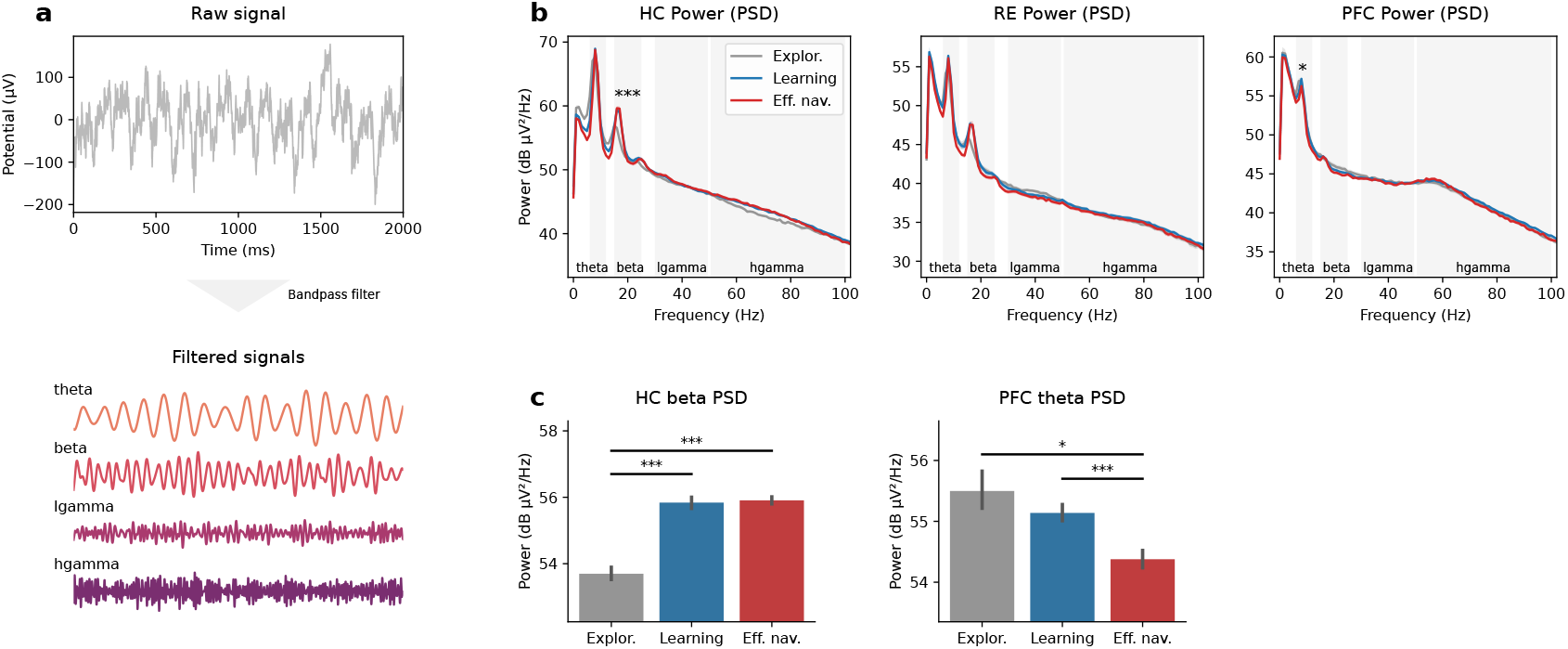
Region-specific changes of the local field potentials (LFP) power spectrum across different learning phases. **(a)** Example of the band-pass LFP filtering. Raw LFP signal was bandpass-filtered into four frequency bands: theta (6-11 Hz), beta (15-25 Hz), low gamma (lgamma, 30-50 Hz), and high gamma (hgamma, 50-100 Hz). **(b)** Power spectral density (PSD) for each recording site (from left to right: HC, RE, PFC). PSD was estimated using Welch’s method with 1s epochs (0.5s overlap). Analysis was restricted to epochs with mean velocity >5cm/s. Conventional band ranges (gray bars) were used, and PSD peaks and troughs. All regions showed elevated theta power; note a second beta peak in the HC and RE power spectra. **(c)** Post-hoc pairwise comparison of HC beta (left) and PFC theta power (right). HC beta power significantly increased from exploration to goal-oriented learning (*T* (5) = 14.34, *p* < 0.001) and remained elevated during efficient navigation (goal-oriented learning vs. efficient navigation: *T* (5) = −0.20, *p* = 0.847). PFC theta decreased towards the end of learning, with significantly lower power during efficient navigation than during goal-oriented learning (*T* (5) = 10.16, *p* < 0.001).

A repeated-measures ANOVA (learning phase × frequency band) revealed significant modulation of HC beta and PFC theta power by learning phase (HC beta: *F* (2, 10) = 81.34, *p* < 0.001, 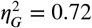; PFC theta: *F* (2, 10) = 8.98, *p* = 0.0351, 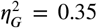); post-hoc pairwise comparisons are summarized in ***Figure 2*** c. HC beta power increased with the onset of goal-oriented learning and remained elevated during efficient navigation. In contrast, PFC theta power gradually decreased, with the lowest values during efficient navigation. No significant learning-phase related power changes were observed in RE.

#### Prominent RE-HC theta-beta coupling emerges with goal-directed navigation

Given the established role of hippocampal theta in entraining higher-frequency oscillations both locally (***Tort et al., 2008, 2009; Shirvalkar et al., 2010***) and across brain regions (***Sirota et al., 2008; Tamura et al., 2017; Hallock et al., 2016***), we next estimated cross-frequency phase-amplitude coupling (PAC) between hippocampal theta phase and higher-frequency amplitudes in RE. In addition, we also quantified HC-PFC PAC, which has previously been reported during spatial learning (***Tamura et al., 2017; Hallock et al., 2016***) was used as a reference.

The strength of PAC was estimated using the modulation index (MI, ***Tort et al. (2008***)), calculated for 10s epochs (5s overlap) and normalized against a surrogate distribution. A repeated-measures ANOVA revealed a significant main effect of learning phase on HC-RE theta-beta PAC (*F* (2, 10) = 24.91, *p* < 0.001, 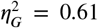). When rats transitioned from exploration to goal-oriented behavior, a prominent PAC peak emerged at ∼8Hz HC theta phase and ∼22Hz RE beta amplitude (***Figure 3*** a), with a preferred phase of 22.86° (***Figure 3*** b). This PAC remained elevated during efficient navigation (***Figure 3*** c).

**Figure 3.**
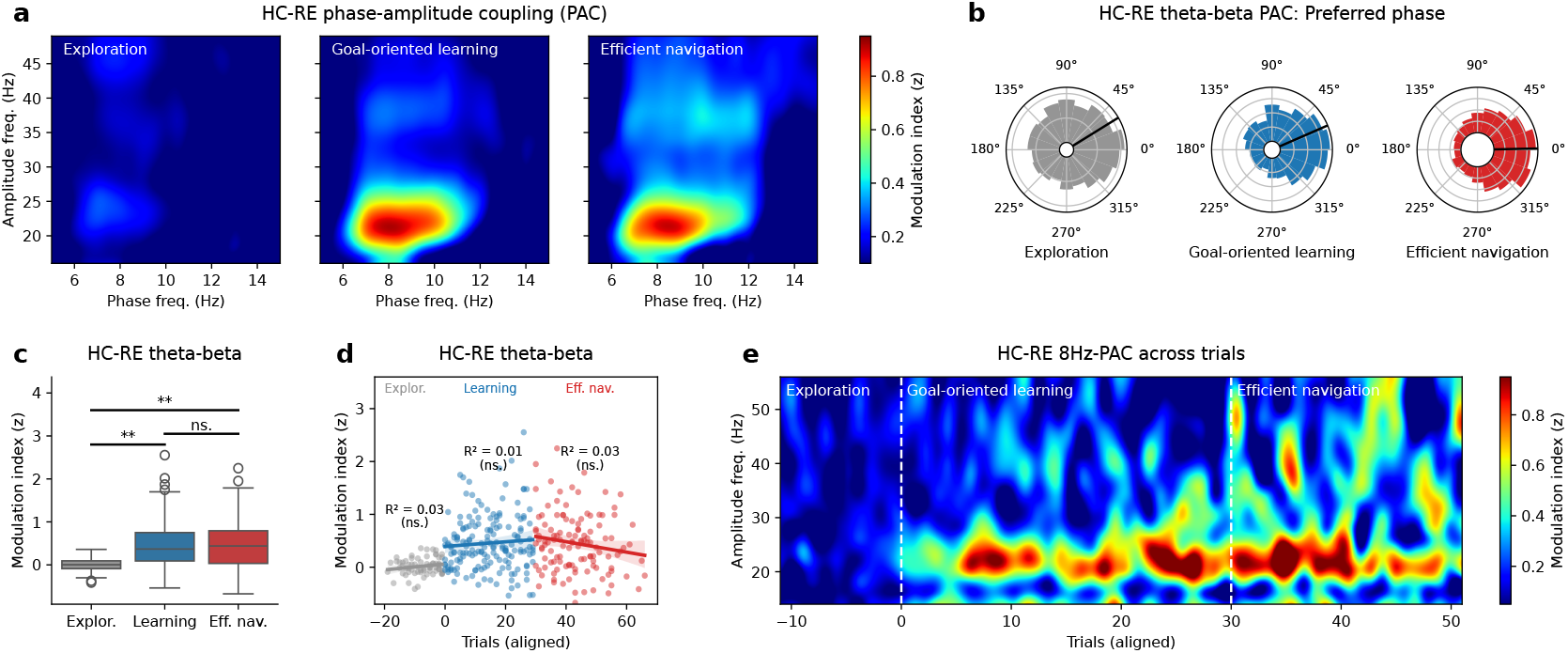
Phase-amplitude coupling (PAC) emerges after the shift in navigational strategy. **(a)** Phase-amplitude comodulogram between the phase of the HC LFP low-frequency band (5-15Hz) and the amplitude of the RE LFP higher-frequency band (16-49Hz), split by learning phase. Note a clear HC-RE theta-beta PAC peak emerging during goal-oriented learning and efficient navigation. The comodulograms also hint at a potential theta-low gamma PAC during efficient navigation, yet the difference compared to the navigation learning phase did not reach statistical significance (*F* (2, 10) = 1.99, 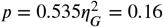). The long temporal windows used in PAC estimation (10s epochs with 5s overlap) may have obscured finer phase-amplitude modulation, particularly at higher frequencies. **(b)** Preferred phase distribution of HC-RE theta-beta PAC, split by learning phase. Black lines indicate the mean preferred phase (Exploration: 31.15°; Goal-oriented learning: 22.86°; Efficient navigation: 1.30°). The lack of significant PAC during the exploration phase is reflected in the wide distribution of preferred phases, which narrowed with the onset of goal-oriented learning. **(c)** Post-hoc pairwise comparison of HC-RE theta-beta PAC. MIs significantly increased from exploration to goal-oriented learning (*T* (5) = −7.89, *p* = 0.002) and remained elevated throughout the efficient navigation phase (goal-oriented learning vs. efficient navigation: *T* (5) = 0.88, *p* = 0.417). **(d)** Linear regression of triwalwise HC-RE theta-beta PAC, split by task phase. While PAC strength significantly increased from exploration to goal-oriented learning (*T* (5) = 7.89, *p* = 0.002), no consistent changes within each learning phase were revealed (exploration: *R*^2^ = 0.03, *p* = 0.171; goal-oriented learning: *R*^2^ = 0.01, *p* = 0.340; efficient navigation: *R*^2^ = 0.03, *p* = 0.083). **(e)** Trial-resolved HC-RE PAC comodulogram, showing 6-11Hz HC theta phase and 14-54Hz RE amplitude coupling per trial, aligned to the exploration-goal oriented learning transition. The comodulogram indicates that theta-beta PAC emerged immediately with the onset of goal-oriented learning.

To assess whether PAC strength changed gradually within learning phases or abruptly at the strategy switch, we fit linear regressions to z-scored trial-wise HC-RE theta-beta modulation indices for each learning phase (***Figure 3*** d). No significant within-phase trends were detected (all *R*^2^ < 0.03, *p* > 0.249), supporting a step-like increase at the transition from exploration to goal-oriented learning (cf. comodulogram in ***Figure 3*** e). This pattern suggests that HC–RE theta–beta coupling reflects a discrete reconfiguration of functional connectivity associated with the behavioral strategy switch, rather than a gradual optimization with ongoing learning.

No significant learning phase effects were observed for HC–RE theta–gamma coupling (theta-low gamma: *F* (2, 10) = 1.51, *p* = 0.535, 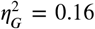; theta-high gamma: *F* (2, 10) = 1.99, *p* = 0.535, 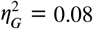). Likewise, in contrast to prior reports linking HC-PFC theta-gamma PAC to spatial working memory (***Tamura et al., 2017; Hallock et al., 2016***), we found no learning-related PAC changes in the HC-PFC pair (main effect of learning phase: all *F* (2, 10) < 0.84, *p* > 0.693, 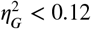).

#### Cross-regional phase synchronization further implicates HC-RE beta

We next examined cross-regional phase synchronization within the HC-RE-PFC network. To this end, we estimated spectral coherence between each pair of regions (HC-RE, HC-PFC, RE-PFC) across learning phases. Coherence was estimated over 1s epochs with 0.5s overlap (***Figure 4*** a).

**Figure 4.**
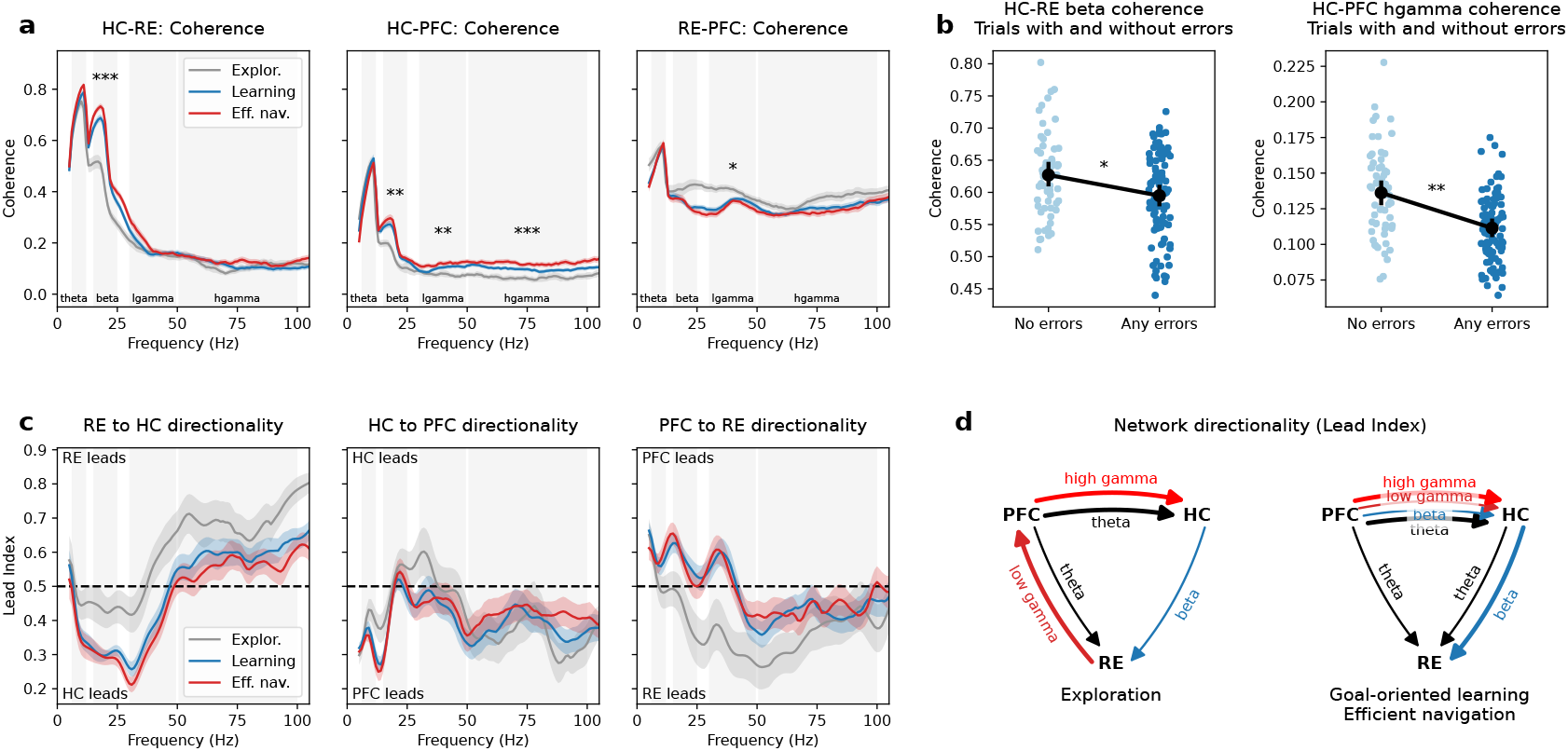
Cross-regional synchrony and direction of information flow during learning. **(a)** Frequency-resolved cross-regional coherence for each region pair, split by learning phase. Gray background bars indicate band ranges for theta, beta, low gamma (lgamma), and high gamma (hgamma). Asterisks indicate significant differences in band coherence (repeated-measures ANOVA, *p* < 0.05: *; *p* < 0.01: **, *p* < 0.001: ***). Line shading shows 95% CIs. All pairs showed elevated theta coherence, which did not vary with learning phase; HC-RE and HC-PFC also peaked in the beta range. Higher frequencies did not show clearly attributable peaks and troughs. **(b)** HC-RE beta and HC-PFC gamma coherence were significantly increased in error-free trials, or, conversely, significantly reduced in trials with errors (HC-PFC beta: *F* (1, 5) = 22.40, *p* = 0.031, 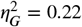; HC-PFC high gamma: *F* (1, 5) = 82.81, *p* = 0.003, 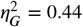). **(c)** Frequency-resolved lead indices (LIs) for each region pair, indicating the direction of information flow. LIs >0.5 indicate that the first region of the pair led the second, while LIs <0.5 indicate that the second region led the first. Gray background bars show band ranges as in (a), and line shading 95% CIs. **(d)** Schematic direction of information flow in the HC-RE-PFC network, split by learning phase. Only connections with significant LIs are shown, as in ***Table 2***. Thick connections signify strong directionality (LI<0.4 or LI>0.6, respectively). Although there were slight changes in lead indices between goal-oriented learning and efficient navigation (***Table 2***), there were no major changes in directionality.

Across all pairs, coherence peaked in the theta band (∼0.5-0.8), with the highest value (0.78) for HC-RE. Theta coherence remained stable across learning phases for all region pairs (all *F* (2, 10) < 1.20, *p* > 0.453, 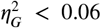). To control for potential overestimation of coherence due to volume conduction, which may be particularly prominent at lower frequencies (***Bastos and Schoffelen, 2016***), we additionally computed the volume-conduction-resistant weighted phase-lag index (***Vinck et al., 2011***). We observed similar theta connectivity patterns (Suppl. ***Figure 1***), suggesting genuine interactions rather than spurious coupling caused by volume conduction.

**Table 2.**
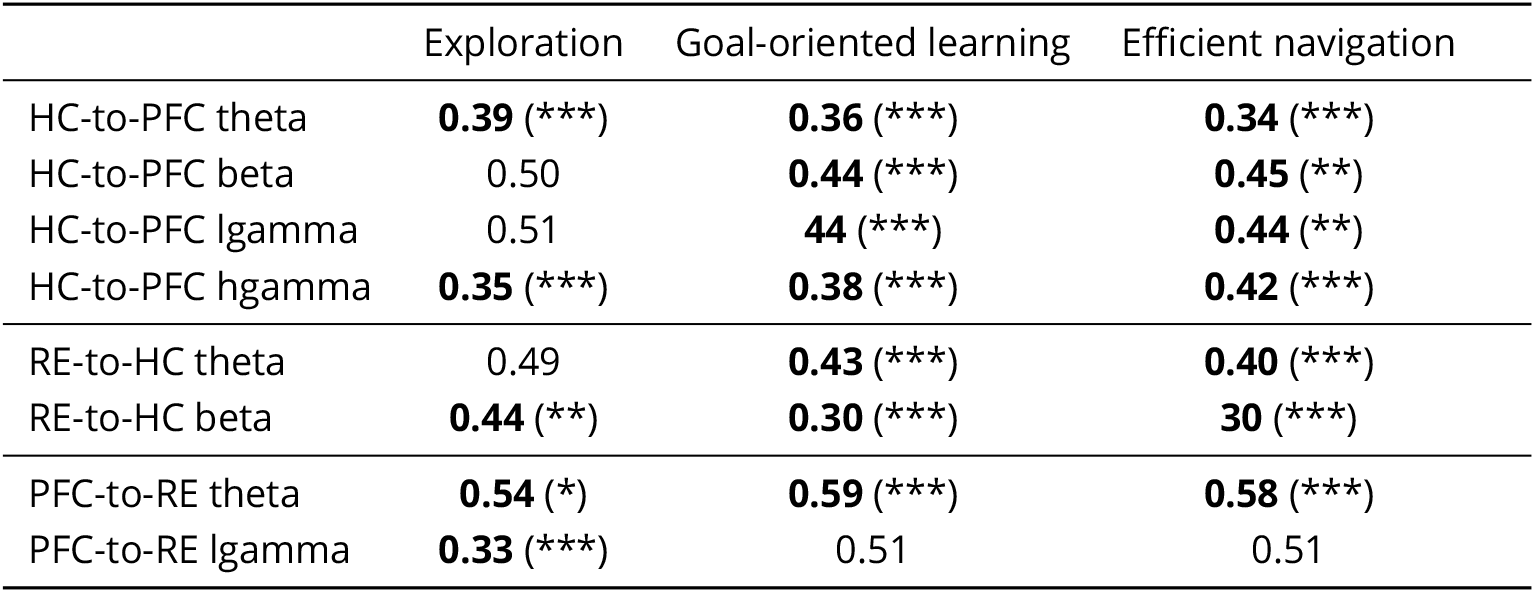
Granger causality lead indices for ordered region pairs (e.g., RE-to-HC), indicating the relative contribution of each region to the total direction of information flow. LIs >0.5 indicate that the first region of the pair led the second, while LIs <0.5 indicate that the second region led the first. Values in bold showed a significant deviation from LI=0.5, as indicated by the asterisks (one-sample t-test, *p* < 0.05: *; *p* < 0.01: **, *p* < 0.001: ***).

|  | Exploration | Goal-oriented learning | Efficient navigation |
| --- | --- | --- | --- |
| HC-to-PFC theta | **0.39 (\*\*\*)** | **0.36 (\*\*\*)** | **0.34 (\*\*\*)** |
| HC-to-PFC beta | 0.50 | **0.44 (\*\*\*)** | **0.45 (\*\*)** |
| HC-to-PFC lgamma | 0.51 | **0.44 (\*\*\*)** | **0.44 (\*\*)** |
| HC-to-PFC hgamma | **0.35 (\*\*\*)** | **0.38 (\*\*\*)** | **0.42 (\*\*\*)** |
| RE-to-HC theta | 0.49 | **0.43 (\*\*\*)** | **0.40 (\*\*\*)** |
| RE-to-HC beta | **0.44 (\*\*)** | **0.30 (\*\*\*)** | **0.30 (\*\*\*)** |
| PFC-to-RE theta | **0.54 (\*)** | **0.59 (\*\*\*)** | **0.58 (\*\*\*)** |
| PFC-to-RE lgamma | **0.33 (\*\*\*)** | 0.51 | 0.51 |

Compared to the other pairs, PFC-RE showed elevated gamma coherence (∼0.4 vs. ∼0.1) throughout the experiment, but varied little with learning. The pair showed a main effect of learning phase on low gamma only (*F* (2, 10) = 7.38, *p* = 0.026, 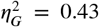, ***Figure 4*** a), driven by a decrease from exploration to efficient navigation (*T* (5) = 3.62, *p* = 0.0456).

For the HC-RE and HC-PFC pairs, the repeated-measures ANOVA revealed significant main effects of learning phase on beta coherence (HC-RE: *F* (2, 10) = 35.63, *p* < 0.001, 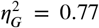 HC-PFC: *F* (2, 10) = 12.92, *p* = 0.005, 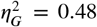, ***Figure 4*** a). In both cases, coherence increased from exploration to goal-oriented learning (HC-RE: *T* (5) = −6.68, *p* = 0.0017; HC-PFC: *T* (5) = −5.00, *p* = 0.0123) and remained thereafter (goal-oriented learning vs. efficient navigation: HC-RE: *T* (5) = −1.83, *p* = 0.1264; HC-PFC: *T* (5) = −0.84, *p* = 0.4381). The similarity in learning-related dynamics between HCRE beta coherence and HC-RE theta-beta PAC suggests that the measures may reflect a common underlying mechanism of cross-regional communication.

Additionally, HC-PFC low gamma (*F* (2, 10) = 19.61, 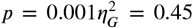) and high gamma coherence (*F* (2, 10) = 33.83, *p* < 0.001, 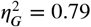) increased progressively across learning.

Previous studies have found links between increased network coherence and working memory performance, in particular for HC-PFC theta and gamma coherence (***Benchenane et al., 2010; Hallock et al., 2016; Jones and Wilson, 2005; de Mooij-van Malsen et al., 2023; Spellman et al., 2015***), but also the HC-RE beta band (***Jayachandran et al., 2023***). Such transient increases tend to occur at maze intersections (***Benchenane et al., 2010; Tamura et al., 2017; de Mooij-van Malsen et al., 2023***) or during the delay period in delay-based working memory tasks (***Jayachandran et al., 2023***) and have therefore been related to decision-making.

Given the sequential decision-making nature of our experiment, we likewise expected differential functional coupling (coherence and/or PAC) depending on navigational efficiency. To test this hypothesis, we selected trials from the efficient navigation phase and split them into sets with (*n* = 94) and without (*n* = 64) errors. HC-PFC high gamma and HC-RE beta coherence were significantly increased in error-free trials (*T* (5) = 9.78, *p* = 0.002 and *T* (5) = 4.93, *p* = 0.026, respectively, fdr-corrected paired t-test; ***Figure 4*** b). We did not observe significant error-based differences in HC-PFC or HC-RE PAC (all |*T* (4) | < 3.32, *p* > 0.176).

#### Directionality of cross-regional communication

We finally examined whether the direction of information flow between regions changed with learning. Granger causality lead indices (LIs) were calculated for ordered region pairs (e.g., RE-to-HC), representing the proportion of total bidirectional information flow attributable to the first region in the pair ((***Hallock et al., 2016***)). LIs > 0.5 indicate a lead of the first region in the pair, and LIs < 0.5 a lead of the second (***Figure 4*** c). Only bands showing significant coherence were analyzed for directionality. Results are summarized in ***Table 2*** and ***Figure 4*** d.

Across all learning phases, the network exhibited significant PFC-to-HC and PFC-to-RE theta directionality. During exploration, RE-to-PFC low gamma directionality and weaker HC-to-RE beta directionality were also present. In line with a decrease of PFC-RE low gamma coherence, the RE-to-PFC low gamma directionality disappeared with the onset of goal-oriented learning. Similarly, HC-to-RE beta directionality increased in parallel with HC-RE beta coherence, and significant HC-to-RE theta directionality emerged. Finally, in addition to persistent PFC-to-HC theta and high gamma information flow, both goal-oriented learning and efficient navigation showed significant PFC-to-HC beta and low gamma directionality.

We did not observe a reversal in information flow, and patterns remained largely stable from goal-oriented learning to efficient navigation. The results further support a functional network reorganization with a switch in navigational strategy, as well as the emergence of HC-RE beta-band communication during goal-directed navigation.

## Discussion

In the present study, we characterized dynamic changes in long-range functional connectivity within the HC–RE–PFC network as rats learned to efficiently navigate a complex maze to obtain a food reward. Learning was associated with a robust increase in beta-band coherence between HC and RE, enhanced HC-RE theta-beta PAC, increased HC beta power, and stronger HC-to-RE Granger causality.

These network changes reflected behavioral performance: error-free trials showed higher HC–RE beta coherence, whereas trials with deviations from the optimal path were accompanied by reduced coupling. Together, our findings provide evidence for a strong hippocampal influence on RE during goal-directed navigation and support an emerging view that RE may act as a working memory buffer for route-related information, forming a potential mechanism underlying flexible spatial behavior.

### HC-RE beta coupling as a spatial memory buffer

Initially recorded in human and non-human primate motor cortex, beta oscillations (∼15-30 Hz) have traditionally been linked to sustained motor activity such as steady muscle contractions (***Baker, 2007***) or the maintenance of the existing motor set (***Gilbertson et al., 2005; Engel and Fries, 2010; Spitzer and Haegens, 2017***). More recent work suggests that beta activity may play a broader role in the active maintenance of information across domains, including working and long-term memory, decision-making, and reward processing (***Lundqvist et al., 2016; Schmidt et al., 2019***).

Our findings extend this role to the spatial domain. HC–RE beta coupling emerged during learning of a maze task requiring sequential navigational decisions, consistent with a recent odor sequence study in rats (***Jayachandran et al., 2023***), where HC-RE and HC-PFC beta coherence correlated with accuracy. The study proposed that transient RE beta activity and HC-RE beta synchrony may serve as a neurobiological “memory buffer” that temporarily maintains task-relevant information and protects it from task-irrelevant interference (***Jayachandran et al., 2023***).

In our study, HC–RE beta coherence was highest in correct, in-sequence trials and reduced for trials with errors, suggesting that HC-RE beta coherence may support the maintenance of a navigational plan across multiple decision points. HC-RE theta-beta PAC and elevated beta coherence during goal-directed behavior, but not during exploration, further suggest that this buffer may be selectively engaged when animals must retain and execute a planned route.

It remains inconclusive whether HC-RE beta coherence and theta-beta PAC reflect a single mechanism or distinct network functions. Compared to beta coherence, we did not observe error-related modulation of theta-beta PAC, but this may reflect a difference in temporal or spectral sensitivity rather than a functional dissociation. Nonetheless, our findings converge with previous work in supporting a specific role for HC–RE beta synchrony in accurate, memory-guided navigation.

Granger causality indicated a dominant HC-to-RE beta directionality. This might appear inconsistent with a buffering role for RE, which could imply RE-to-HC signaling. However, we argue that a hippocampal drive to RE is compatible with a model where RE updates and maintains sequence representations, relaying them back to HC (or PFC) as needed. Anatomical evidence supports this bidirectional connectivity, showing reciprocal connections between RE and ventral CA1 (***Varela et al., 2014; Dolleman-van der Weel et al., 2019; Cassel et al., 2013, 2021***).

Lesion studies also align with this interpretation: RE is essential for tasks requiring spatial memory (***Davoodi et al., 2009; Mei et al., 2018; Hembrook and Mair, 2011***), sequence memory (***Jayachandran et al., 2019***), object recognition (***Barker and Warburton, 2018***), and fear memory acquisition (***Xu and Südhof, 2013***). On the other hand, lesioning RE shows no effect in tasks that do not require HC-PFC communication, such as those depending on HC (***Dolleman-van der Weel et al., 2009; Hembrook et al., 2012; Loureiro et al., 2012; Cassel et al., 2013***) or PFC only (***Ito et al., 2015***), and has no effect on post-trial memory consolidation (***Mei et al., 2018***). The pattern suggests a specific role for RE in online memory coordination. Interestingly, despite pronounced behavioral shifts with learning, RE power spectra were stable, and changes mainly emerged in cross-regional beta coherence, suggesting that learning involves a dynamic reorganization of network interactions rather than changes in local RE activity.

We also observed PFC-RE low gamma modulation: coherence was high during exploration but decreased with the onset of goal-oriented learning. Although we observed primarily RE-to-HC directionality, the results may indicate a shift in dominant RE inputs from PFC during exploration to HC during goal-directed navigation, consistent with recent work suggesting a specific role of PFC-RE communication in non-HC-dependent memory (***Ciacciarelli et al., 2025***).

### Hippocampal-prefrontal gamma coherence and efficient navigation

PFC-to-HC gamma coherence, though initially low, increased progressively with learning. Both low and high gamma coherence followed behavioral performance more closely than other bands, increasing with navigational efficiency and distinguishing between all three learning phases. The findings align with previous studies linking HC-PFC gamma-band synchrony to working memory and decision-making (***Bygrave et al., 2019; de Mooij-van Malsen et al., 2023***). Velocity differences between learning phases are unlikely to fully explain these effects, as previous work showed that PFC-HC gamma coherence peak frequency, but not magnitude, is modulated by running speed (***Ahmed and Mehta, 2012***). However, because accuracy and speed covaried in our task, we cannot completely disentangle cognitive from motor contributions.

### Theta coordination remained stable across learning

HC-PFC theta synchrony has previously been linked to spatial working memory and decision-making, showing increased coherence for correct decisions in working-memory tasks (***Jones and Wilson, 2005; Hyman et al., 2010; Hallock et al., 2016; de Mooij-van Malsen et al., 2023***). We also observed strong theta coherence across all region pairs, but it did not vary with learning phase or trial accuracy. Granger causality consistently showed PFC-to-HC-to-RE theta directionality, but this pattern was likewise stable and unrelated to performance.

These results argue against dynamic, behavior-dependent theta coordination within the HC-REPFC network during goal-directed navigation. Rather, theta synchrony may reflect a baseline state of anatomical connectivity between regions, whereas beta and gamma interactions are more sensitive to task demands. However, we note that the relatively coarse temporal scale in our analyses may have masked transient theta dynamics seen in single-choice paradigms. Follow-up eventaligned analyses for individual decision points may reveal more subtle modulations.

### Limitations and future directions

Several limitations should be acknowledged. Firstly, spatial working memory was not directly assessed in the present study, although it was likely engaged, as efficient navigation of the maze required tracking of the current position and prior choices. However, we cannot fully rule out that the task may have recruited multiple memory systems, including procedural learning (e.g., rote memorization of a sequence of turns), particularly towards the end of learning. However, procedural learning is unlikely to depend on RE, as action sequence learning is unaffected in RE-lesioned rats (***Hembrook and Mair, 2011***).

Secondly, our LFP-based approach, while informative about network-level dynamics, cannot resolve cellular mechanisms or distinguish between genuine cross-regional interactions and common inputs from a third region. Future studies combining single-cell recordings with pathwayspecific manipulations will be essential for establishing causal relationships.

Lastly, trial-averaged analyses may have obscured transient dynamics accompanying specific behaviors, for example, at choice points or during error correction. Event-locked, high-resolution analyses are required to reveal more precise patterns of network coordination.

## Methods and Materials

### Animals

Experiments were conducted in adult male Sprague-Dawley rats (n = 6; Charles River Laboratories, Sulzfeld, Germany) weighing 300-350g at the start of the experiment. The initial cohort consisted of 17 rats, of which six rats had confirmed electrode placement in all target regions (PFC, HC, and RE).

Rats were housed in a controlled facility maintained at 20-30°C temperature, 40-60% humidity, and a 12h/12h light/dark cycle. All experiments were performed during the dark cycle. Before chronic electrode implantation, rats were group-housed with ad libitum access to food and water. Following surgery, rats were single-housed to prevent implant damage. During behavioral testing, rats were food-restricted to 90% of ad-libitum body weight to ensure their appetitive motivation.

All procedures conformed to Directive 2010/63/EU of the European Parliament and the German Animal Welfare Act (TierSchG and TierSchVersV). Protocols were approved by the local ethics committee (§15 TierSchG) and the state authority (Regierungspräsidium Tübingen, Baden-Württemberg, Referat 35, Veterinärwesen).

### Surgical procedures

All surgeries were performed under standard aseptic conditions. Rats were anesthetized with isoflurane (5% induction, 1.5-2% maintenance) and head-fixed in a stereotaxic frame (David Kopf Instruments, Tujunga, CA) with head angle adjusted to 0°. Body temperature (∼37°C), heart rate, and blood oxygenation (above 90%) were continuously monitored.

After subcutaneous lidocaine (Lidocard 2%, B. Braun, Melsungen, Germany), the skull was exposed and three craniotomies were performed above the right-hemisphere target regions according to the rat brain atlas (***Paxinos and Watson, 2006***): PFC (AP/ML = 3.1/0.8mm), HC (AP/ML = - 3.8/2.4mm), and RE (AP/ML = -1.8/-1.3mm). Single-tipped platinum-iridium electrodes (FHC Inc., Bowdoin, ME) were chronically implanted at the following coordinates: PFC (DV = 3.4mm), HC (DV = 2mm), and RE (DV = 6.8mm, 10° medial-lateral angle to avoid the sagittal sinus).

Five stainless steel screws (0.86mm diameter) were fixed to the skull and served as ground and anchors for the implant. A 3D-printed plastic frame and copper mesh provided electrical shielding and were secured to the skull with tissue adhesive. The entire implant was fixed with dental cement (RelyX Unicem 2 Automix, 3M, MN).

During post-surgery recovery (5-7 days), rats received antibiotics (Baytril 5mg/kg i.p., Bayer) and analgesics (Rymadil 2.5 mg/kg i.p., Zoetis).

### Behavioral apparatus and training

Behavioral testing took place in a custom-built “crossword maze” (130 × 130cm, black PVC) composed of 4 × 4 arms arranged in a lattice, elevated 80cm above the floor. Maze arms were 10cm wide with 2cm rims. A reward port (lower left corner, ***Figure 1*** a) dispensed chocolate milk (0.6ml) upon nose poking. Black curtains surrounded the maze, and experiments were conducted under dim light. Two large posters affixed to the curtains served as distal visual cues.

Before the experiment, rats were habituated to the experimental environment over 1 to 3 sessions. The food-deprived rats were released into a free-roam version of the maze without blocked arms and allowed to consume chocolate milk distributed throughout the maze. The habituation period concluded when the rats readily consumed the chocolate milk and showed no signs of anxiety.

For the experiment, nine vertical barriers (30cm high, 25-40cm wide) were inserted to block specific arms (***Figure 1*** a). At the beginning of each trial, rats were released from one of two pseudorandomized starts (A1 or B1, ***Figure 1*** a) and had up to 3 minutes to reach the reward port.

Trials ended upon reward consumption or after 3 minutes. Between trials, rats rested in a waiting box (3-5 min) while the maze was cleaned to remove olfactory cues. Each rat completed 10 trials per session for seven consecutive days; one rat had a session with only eight trials.

### Behavioral tracking

Movement was recorded at 25Hz using an overhead camera that tracked two head-mounted LEDs, time-stamped in sync with the electrophysiological recording system. The head position was estimated by averaging LED pixel coordinates. Tracking dropouts due to obstruction were linearly interpolated. Velocity and orientation were derived from smoothed positional derivatives (Savitzky-Golay filter). Body pose and orientation were not tracked.

Task performance was assessed using: (1) trial duration as time from start to reward, (2) mean velocity, (3) time spent immobile (velocity <5cm/s), and (4) navigational efficiency (error proportion relative to total navigational decisions, see below).

The maze was divided into 16 zones defined by 20cm radius circles around each intersection (***Figure 1*** a). Using these zones, trajectories were partitioned into discrete events; events <0.12s (3 frames) were discarded to avoid aliasing effects at zone boundaries. Optimal paths (A1-C6 or B1-C6) were defined, and trajectories were compared to the optimal paths. At any given intersection, a navigational decision could either decrease or increase the distance towards the goal. The former was counted as correct, while the latter was counted as an error. The maze contained no loops, ensuring that there was always a correct choice.

Navigational efficiency was calculated as the fraction of incorrect choices among all valid decisions. Only intersections offering more than one choice were included.

### Recording procedures and histology

After post-surgical recovery, rats were habituated to moving freely while connected to the recording cable. The implants were connected to a Neuralynx Digital Lynx acquisition system (Neuralynx, Bozeman, MT) through a 32-channel headstage (Neuralynx, Bozeman, MT) and a custom adapter (SSD-10-SS-GS, Omnetic, Minneapolis, MN). Broadband signals (1-9000Hz) were digitized at 32kHz.

Following the final recording session, rats were euthanized (Narcoren, 100mg/kg i.p., Merial) and transcardially perfused with 0.9% saline followed by 4% paraformaldehyde. Brains were sectioned horizontally (50 µm; Microm HM 440E, Thermo Fischer Scientific, Waltham, MA), and electrode placements were manually verified in Nissl-stained sections.

### Signal processing

Raw LFPs were downsampled from 32 kHz to 1 kHz after applying a zero-phase-delay FIR antialiasing low-pass filter at the Nyquist frequency. Segments (∼0.5s) with amplitudes >±2mV were automatically excluded, followed by manual inspection. Trials with >50% rejected data were removed (3 trials, 0.72%). One full session (10 trials, 2.39%) was excluded due to persistent broadband noise.

Signals were filtered into discrete frequency bands using the following band ranges: theta (6-11Hz), beta (15-25Hz), low gamma (30-50Hz), and high gamma (50-100Hz).

### Quantification and statistical analysis

Analyses were performed in Python 3 using custom scripts with Scipy, MNE, MNE-connectivity, and tensorpac. Data were aggregated per trial, then averaged per animal and learning phase. For temporal analyses, trials were aligned to the last unsuccessful trial per rat, i.e., the last trial in which the rat failed to reach the reward.

Power and coherence were computed in 1s windows with 0.5s overlap (Welch’s method). Only signal windows with clear locomotive behavior (>5cm/s velocity) were included in the computation. Phase-amplitude coupling (PAC) was quantified using the modulation index (MI; ***Tort et al. (2008***)), testing HC theta phase against RE and PFC beta/gamma amplitudes. The magnitude and robustness of the MI depend on the number of cycles per sample, i.e., the length of the compared signal. Following recommendations in ***Hülsemann et al. (2019***), we chose longer temporal windows for PAC estimation (10s epochs with 5s overlap) to obtain robust estimates. PAC was only calculated for epochs with mean velocity >5cm/s. Trials with total duration <10s were also excluded from PAC estimation (goal-oriented learning: 7 trials, 3.89%; efficient navigation: 36 trials, 21.30%). Finally, PAC estimates were z-scored against a surrogate PAC distribution (n=200) to obtain a normalized measure.

Directionality was assessed via Granger causality, following the description in ***Hallock et al. (2016***). For each region pair *a, b*, frequency-resolved Granger causality was computed bidirectionally over 1s windows with 0.5s overlap. Lead indices (LI) were defined as *a* → *b*/(*a* → *b* +*b* → *a*), with LI >0.5 indicating a lead by region *a*. LI significance was tested using one-sample t-tests against 0.5.

In general, statistical significance was assessed using repeated-measures ANOVAs or paired t-tests, with p-values corrected for multiple comparisons using Benjamini-Hochberg FDR. ANOVA result effect sizes were estimated using the generalized Eta square 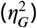 measure. Changes over time were estimated using linear regression. Significance threshold was set at *p* = 0.05.

## Acknowledgments

This work was supported by the Max Planck Society. We thank Peter Dayan, Ivan de Araujo, and Vinod Kumar for their theoretical input and assistance, and Axel Oeltermann and Joachim Werner for their technical support.

## Supplementary material

**Supplementary Figure 1.**
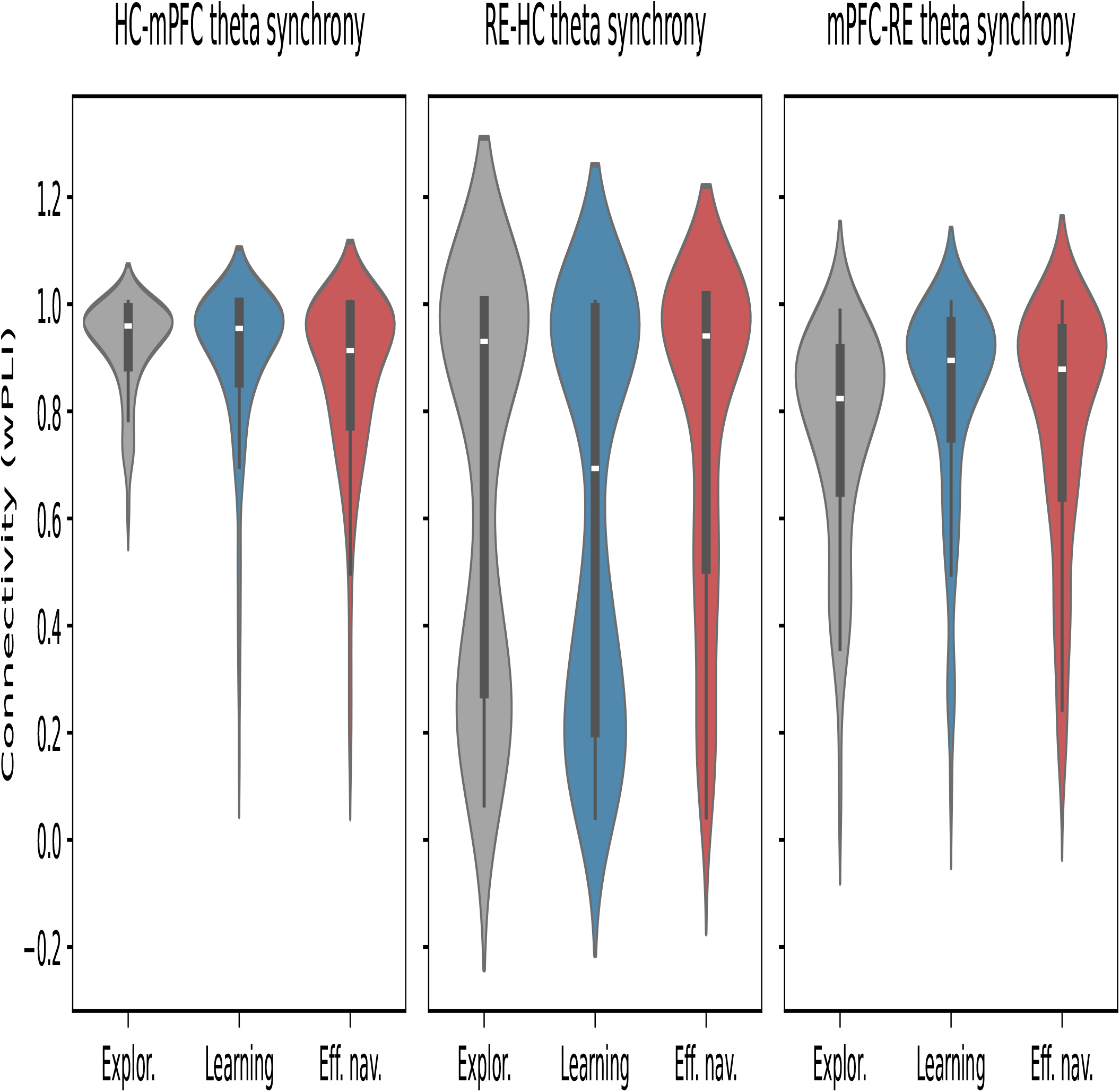
Cross-regional theta synchrony (weighted phase-lag index), split by learning phase.

