## Supplementary figures and images for "Hippocampus-reuniens beta coupling supports goal-directed spatial navigation"

### Supplementary Figure 1

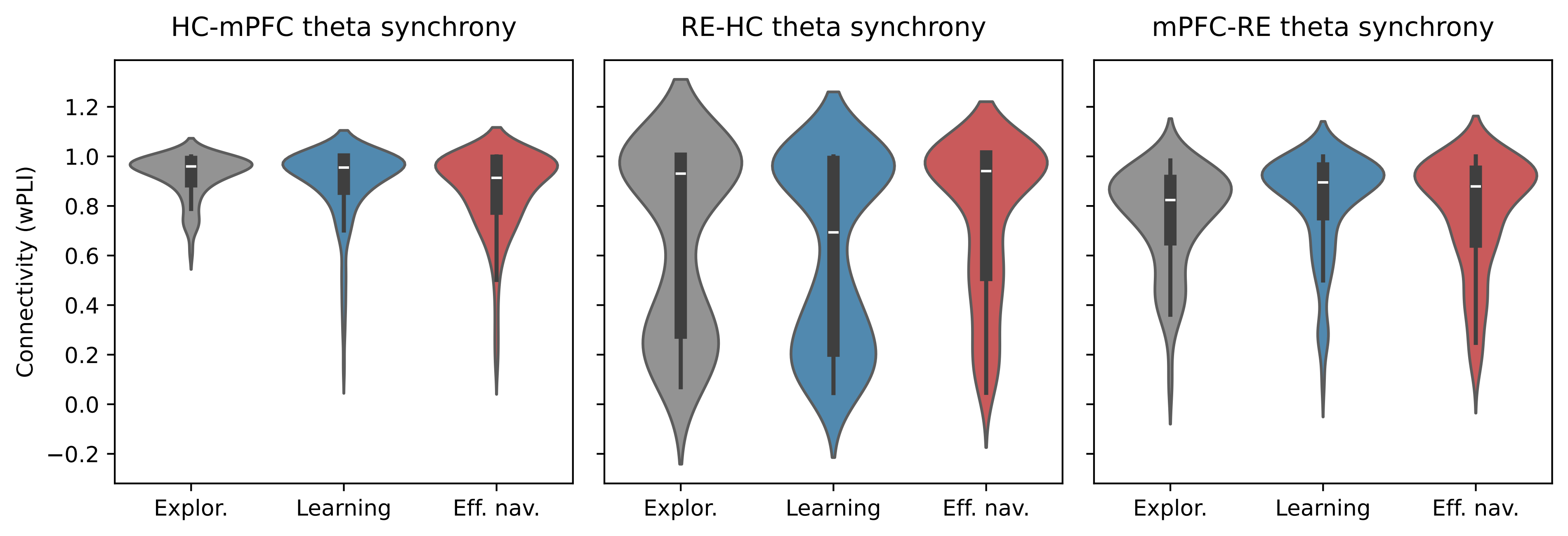
